# A Draft Genome and High-Density Genetic Map of European Hazelnut (*Corylus avellana* L.)

**DOI:** 10.1101/469015

**Authors:** Erik R. Rowley, Robert VanBuren, Doug W. Bryant, Henry D. Priest, Shawn A. Mehlenbacher, Todd C. Mockler

## Abstract

European hazelnut (*Corylus avellana* L.) is of global agricultural and economic significance, with genetic diversity existing in hundreds of accessions. Breeding efforts have focused on maximizing nut yield and quality and reducing susceptibility to diseases such as Eastern filbert blight (EFB). Here we present the first sequenced genome among the order Fagales, the EFB-resistant diploid hazelnut accession ‘Jefferson’ (OSU 703.007). We assembled the highly heterozygous hazelnut genome using an Illumina only approach and the final assembly has a scaffold N50 of 21.5kb. We captured approximately 91 percent (345 Mb) of the flow-cytometry-determined genome size and identified 34,910 putative gene loci. In addition, we identified over 2 million polymorphisms across seven diverse hazelnut accessions and characterized t heir effect on coding sequences. We produced t wo high-density genetic maps with 3,209 markers from an F1 hazelnut population, representing a five-fold increase in marker density over previous maps. These genomic resources will aide in the discovery of molecular markers linked to genes of interest for hazelnut breeding efforts, and are available to the community at https://www.cavellanagenomeportal.com/.

## Introduction

European hazelnut (*Corylus avellana*, L.) is a member of the family Betulacea, a group of six plant genera that include the birches (*Betula* L. spp.) and alders (*Alnus* Mill spp.). Members of this family are either a reduced stature tree or are shrub-like with fruit produced in the form of kernels. It is these kernels, the hazelnuts, which make *C. avellana* an agriculturally significant crop. Hazelnuts provide the predominant flavor in a variety of butters, candies, chocolate spreads, and confectionary pastes. Whole blanched and roasted kernels are in demand by consumers worldwide, and the shells are used for both landscaping and groundcover. Hazelnuts are also high in fiber, contain several essential vitamins, and have potential for use as biofuels (1).

European hazelnut grows best in areas with mild coastal tempered winters, and the vast majority of the world’s hazelnut production is centered in the Black Sea region of Turkey. This region accounts for over 70 percent of hazelnuts on the global market (2), producing over 1 million tons each year. The similarly moderate climate of Oregon’s (USA) Willamette Valley is also well suited for hazelnut production. Although Oregon provides for only 3 perrcent of the world share of hazelnuts, this region produces 99 percent of the hazelnuts grown for the North American market worth an estimated US 67.5 million dollars annually, according to the USDA NASS Oregon Field Office, 2011.

Hazelnut is clonally propagated using traditional simple layerage, tie-off layerage (stooling), or grafting, all of which are labor-intensive and produce limited numbers of plants (3). Micropropagation, or in vitro propagation on a defined culture medium, allows rapid propagation of selected types, and has been particularly useful in the rapid increase of new cultivars and pollinizers. Meristems from these plantlets may be subcultured for up to 6 years without change to their genetic structure (3), and storage at 4C allows maintenance of many accessions.

Breeding efforts have focused on producing progeny with enhanced agronomic traits of interest such as increased kernel oil content and size, blanching ability, bud mite resistance, and disease resistance. Specifically, development of germplasms resistant to the fungus Anisogramma anomola (Peck) E. Muller, the causal agent of Eastern filbert blight (EFB), is important. EFB is a deadly disease that initially causes cankers to form on the woody parts of the tree. These cankers act to slowly girdle the branches, leading to canopy leaf loss and death of the tree within a few years (4). Most of the European hazelnut varieties grown in North America are susceptible to EFB including ‘Barcelona’, the predominant heirloom cultivar grown in Oregon and the Pacific Northwest.

The United States Department of Agriculture (USDA) and Oregon State University (OSU) have collected over 700 accessions of *C. avellana* L., preserved in Corvallis, Oregon where they are used in breeding efforts (5). These accessions are maintained both as plantlets in tissue culture and as adult trees grown near the Oregon State University campus.

Among the European hazelnut accessions preserved in Corvallis is an F1 mapping population of 138 individuals resulting from the cross of the EFB-resistant accession OSU 414.062 (paternal) and EFB-susceptible accession OSU 252.146 (maternal). Among these progeny is the accession ‘Jefferson’ (OSU 703.007), which was released in 2009 by Dr. Shawn Mehlenbacher at Oregon State University(6). ‘Jefferson’ was selected as the reference hazelnut accession for genome assembly and characterization due to the presence of the dominant ‘Gasaway’ allele, which confers resistance to EFB. Identifying sources of resistance and resistant cultivars will not only increase the parental germplasm available for future breeding efforts but will also the reduce fungicide requirements and EFB-associated costs to regional growers (7).

*C. avellana* possesses several traits that make it an attractive candidate for use as a model system for the family Betulacea. Among these are a short life cycle; *C. avellana* seedlings produce fruit 5 years post-planting. It occupies a small habit for a tree at 5 m and is amenable to *Agrobacterium-mediated* transformation through tissue culture methods. European hazelnut is a diploid with 11 chromosomes (2n =2x=22) and a genome of 378 Mbp (empirically determined by flow cytometry at the USDA-ARS-NCGR in Oregon in 2012). This genome is approximately triple the size of the established model dicot *Arabidopsis thaliana* (L.) Heynh, at 125 Mbp and slightly larger than peach at 230 Mbp(8) but is significantly smaller than those of other sequenced tree genomes such as *Populus trichocarpa* at 520 Mbp (9) and loblolly pine at 23 Gbp (10).

We sequenced the ‘Jefferson’ genome at 93x coverage and have completed a de novo genome assembly, capturing 91 percent of the genome (345 Mbp) with a contig N50 of 21,540 bp. Homology-based functional annotation, restricted to whole gene models and aided by the ‘Jefferson’ transcriptome assembly (11) predicted 34,910 protein coding loci. Of these predicted loci, 22,474 have homology to an entry in the NCBI non-redundant protein database, and 82.5 percent of the annotated genes are presented in the best annotated and closest related genera *Vitus, Prunus, Populus*, and *Ricinus*.

Additionally we resequenced seven European hazelnut accessions at ~20x coverage and discovered millions of polymorphisms between one of more of these genomes and that of ‘Jefferson’. We predicted the effects of each polymorphism on the coding potential of affected loci and identified candidate genes for future research. Genotyping by sequence (GBS) analysis of the F1 mapping population enabled identification of 3,209 additional GBS-derived markers between the maternal (OSU 252.146) and paternal (OSU 414.062) maps. This improved high-density genetic map will be useful for marker-assisted breeding and for the identification of new, desirable traits in hazelnut. Hazelnut genome sequencing has provided new resources to the scientific community and promises to accelerate trait discovery and enhance future breeding efforts. This resource will serve as a tool for gene discovery and functional studies, for the development of DNA markers and other genomic tools, and will allow future integration of the genome sequence with genetic and physical maps and the incorporation of new sequencing technologies.

## Materials and methods

### Collection of Tissues and Sequencing on the Illumina HiSeq 2000

Whole leaf tissues were collected from field-grown trees located at the Smith Horticultural Research farm in Corvallis, Oregon and flash frozen in liquid N2. Tissues were ground to a powder in liquid N2 using a mortar and pestle and genomic DNA (gDNA) was extracted using Qiagen Plant DNeasy kits per the manufacturer’s directions. The integrity and quality of the gDNA was assessed by visualization on 1 percent agarose gels and quantified using the Qubit Fluorometer (Life Technologies) prior to library construction. The construction of 250-bp and 350-bp Illumina paired-end (PE) libraries and a 4.5-Kb mate-pair (MP) library was performed at the Georgia Genomics Facility at the University of Georgia. Cluster generation on the Illumina HiSeq 2000 was performed in the Oregon State University Center for Genome Research and Biocomputing (CGRB) core facility using a standard Illumina protocol.

### Genome Assembly and Filtering

Genomic data from two PE libraries (250-bp insert, 350-bp insert) and one MP library (4.5-Kb insert) were trimmed based on an assigned Illumina quality score of Q30 (1 in 1000 change of incorrect base call) and below using the FASTX toolkit version 0.0.13 and 5’ barcode and adaptor sequences were removed. The *de novo* genome assembly package Velvet v1.2.01 (12) was used to generate assemblies of these quality-filtered genomic data with a k-mer hash length of 51 bp. The program SSPACE v2.0 (13) was then applied, using all quality-controlled data, to improve the Velvet assembly by merging and extending scaffolds where possible. Contigs shorter than 1 Kb were discarded, as were those having greater than 25 percent homology and greater than 10 unique best BLASTN hits to annotated non-plant and organelle sequences in the NCBI nucleotide database to remove any bacterial or fungal contamination.

### Gene Prediction and Functional Annotation

Low complexity and repetitive regions in filtered contigs were masked using the program RepeatMasker (14), and putative loci were identified using the ab initio gene prediction software AUGUSTUS (15). We trained AUGUSTUS with the well characterized gene features of *Arabidopsis thaliana* (16). Gene predictions were restricted to whole models only in order to avoid partial gene calls resulting from alternative splicing, incomplete transcripts, and incomplete genes resulting from mis-assembly. The 28,255 putative transcript models from the European hazelnut transcriptome assembly (11) were used by the program as empirical evidence to guide the gene predictions, by offering locational information of gene features. Annotation of the putative amino acid sequences was conducted via sequence homology to the NCBI non-redundant (nr) protein database. The AUGUSTUS output was aligned the to the nr database using BLASTP tool (17) with an E-value cut-off of 1*e*^−10^ resulting in 34,912 putative protein coding regions. After annotation, further filtering was implemented to remove all loci annotated as transposable elements, retroelements, and gag-polymerases via homology to the NCBI non-redundant protein database using the BLASTP tool(17) with an E-value cut-off of 1*e*^−10^. C.avellana Jefferson 1 was _ discarded post-filtering. Additionally, we used the program BLAST2GO to functionally annotate the filtered set of putative amino acid sequences f rom the AUGUSTUS output using an expected value cutoff of 1*e*^−10^ resulting in 65,536 GO terms classifying 11,221 unique loci.

The loci removed from the annotated protein file are available for visualization on the hazelnut JBrowse portal and download on the FTP server hosted at (https://www.cavellanagenomeportal.com).

### Polymorphism Discovery

We sequenced seven unique hazelnut cultivars (‘Barcelona’, ‘Tonda Gentile delle Langhe’, ‘Tonda di Giffoni’,‘Ratoli’, ‘Daviana’, ‘Halls Giant’, ‘Tombul (Extra Ghiaghli)’), uniquely barcoded and pooled in equal molar r atios, on f our lanes of an Illumina HiSeq 2000 flowcell. The resulting reads were quality filtered as described above. The Burrows-’Wheeler Aligner (BWA) was used to align the filtered reads from each of the seven cultivars to the ‘ Jefferson’ reference genome, using default settings (18). We then applied the Genome Analysis Toolkit (GATK) (19) f or base quality score recalibration, insertion and deletion (INDEL) realignment and dupli-cate removal. Single-nucleotide polymorphisms (SNP) and INDEL discovery was performed across all samples simultaneously using standard hard filtering parameters according to GATK Efest Practices recommendations (19,20). In order to increase confidence in polymorphism calls, SNP and INDEL predictions were f urther filtered using the SnpSift component of the SnpEff package (21); a minimum of 15 overlapping reads and a quality score (Q) of 30 f or the SNPs and 20 f or the INDELS were required. SnpEff was used f or genome-wide variant annotation and to predict the effects of each polymorphism on the coding potential of the putative loci.

### Visualization of Data

For visualization of the gene features and polymorphisms we uploaded the alignments, predicted AUGUSTUS gene models, and their BLASTP annotations to JBrowse (http://jbrowse.org/), a Java-based genome browser. These datasets are available for visualization on the hazelnut genome website hosted at https://www.cavellanagenomeportal.com

### Construction of the GBS-based Genetic Map

The high-density genetic map was constructed from a full-sib population (138 seedlings and two parents in triplicate) from the cross of OSU 252.146 and OSU 414.062 using a two-way pseudo testcross approach (22). High-quality genomic DNA from the progeny and parental plants was extracted as previously described and used for construction of GBS libraries (23). GBS libraries were prepared using the restriction enzyme ApeK1 and pooled in sets of 72 uniquely barcoded individuals. Each 72 individual barcoded pool was sequenced (4 lanes total) on an Illumina HiSeq 2500 1×100 SE run. Polymorphisms were identified by implementing the UN-EAK package of the TASSLE-GBS pipeline (24) using default settings. A minimum coverage of 5 overlapping reads was required to call each polymorphism in order to minimize false positives.

A total of eight F1 individuals were removed prior to analysis because of low coverage. Raw genotype output from TASSEL was first filtered to remove SNPs with more than 20 percent missing data in the population. After filtering, SNPs that were homozygous in the maternal parent (OSU 252.146) but heterozygous in the paternal parent (OSU 414.062) and heterozygous in the maternal parent (OSU 252.146) but homozygous in the paternal parent (OSU 414.062) were used f or map construction. As these configurations are expected to segregate at a 1:1 ratio, any SNPs that f ailed to meet this segregation pattern were discarded. The remaining 2,198 SNP markers were converted into a cross pollinator (CP) population and then mapped using JOINMAP 4.1(25). Markers were assigned to linkage groups (LGs) with independence LOD scores of 8.0. After classifying markers into linkage groups, the regression mapping algorithm and a maximum recombination fr equency of 0.40 were employed. Genetic distances between loci were calculated with Kosambi’s function. Homologous regions in peach were identified using BLAST with a minimum e value of 1×10^−5^ and minimum length of 40 bp. Markers mapping with multiple high confidence hits were removed as these likely represented repetitive sequences in peach or duplicated chromosomal regions.

### Data Availability

The current assembly for the ‘Jefferson’ genome is available for BLAST queries and downloads at https://www.cavellanagenomeportal.com/: putative amino acid sequences, variant calls, and effect predictions for the seven resequenced accessions, and the current genetic linkage map are also available for download via FTP. Polymorphism and gene feature tracks of the seven re sequenced hazelnut accessions and ‘Jefferson’, along with functional annotation and variant effect predictions, can be visualized at the European hazelnut web portal.

## Results and Discussion

### Assembly and Characterization of the European Hazelnut Genome

The objective of this study was to provide hazelnut genomic resources for breeders for use in gene and marker discovery and polymorphism detection with the ultimate goal of hazelnut improvement. We assembled the European hazelnut accession ‘Jefferson’ using two PE and one MP Illumina library collectively representing 93x genome coverage with the assembly programs Velvet and SSPACE. We discarded all contigs shorter than 1 Kb, and implemented a nucleotide-filtering cutoff to remove non-plant and organelle sequences from the assembly. After filtering, the draft assembly for the hazelnut genome included 36,641 contigs and scaffolds with a total sequence length of 345 Mb. This is 91 percent of the size of the genome determined by flow cytometry, for approximately 93x coverage. Half of the assembly is contained in scaffolds and contigs greater than 21.5 Kb, with the largest scaffold comprising 274.5 Kb (Table 1).

**Table 1.** Summary of hazelnut genome assembly and annotation

|  |  |
| --- | --- |
| Total number of contigs | 36,641 |
| Total size of assembly | 345.5 Mb |
| Length of largest contig | 274.5 Kb |
| Average contig size | 9.4 Kb |
| N50 | 21.5 Kb |
| Number of *ab initio* predicted protein-coding loci | 34,910 |
| Number of functionally annotated gene products | 22,474 |

The fragmented nature of the assembly is likely to high within genome heterozygosity, evidenced by the bimodal K-mer distribution (Fig 1).

**Fig 1.**
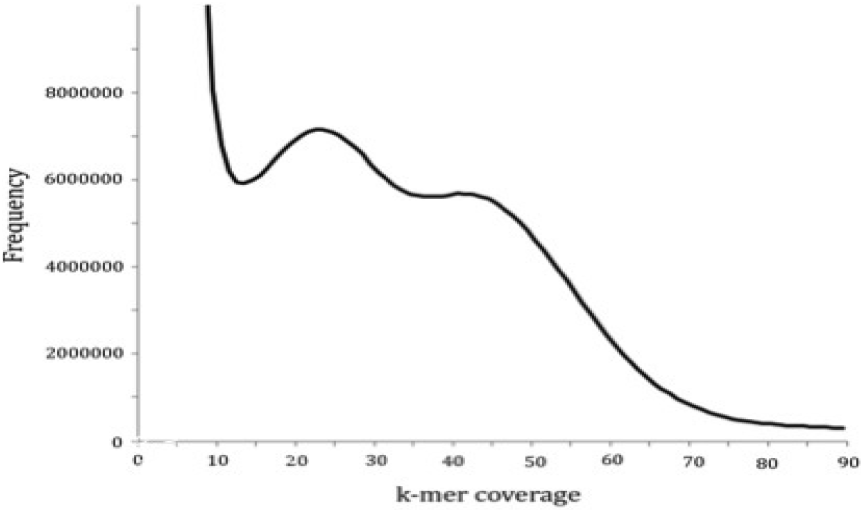
K-mer coverage of the ‘Jefferson’ 350bp PE Illumina library (k=21). The peak with k-mer coverage of 22 represents heterozygous sites and the peak at coverage 44 represents homozygous sites. The large peak at 22X coverage suggests high within-genome heterozygosity.

Marker development in hazelnut has produced roughly 500 random amplified polymorphic DNA (RAPD) markers and 700 simple sequence repeat (SSR) markers (22). We have now constructed a high-density GBS-based map using a full-sib population of 136 F1 plants from the cross of OSU 252.146 and OSU 414.146. The maternal parent, OSU 252.146, is susceptible (S) to EFB, and the paternal parent, OSU 414.062, displays resistance (R) to blight. A total of 21,379 high-confidence SNPs were genotyped across the 136 offspring including 2,198 heterozygous SNPs in OSU 252.146 and 1,966 SNPs in OSU 414.146. Maps were constructed using JoinMap 4.1 with an independence LOD score of 10.0. The maternal genetic map (OSU 252.146) has 1,741 GBS markers and 270 SSR/RAPD markers spanning 762 cM across 11 linkage groups (**Table 2A**).

**Table 2A.**
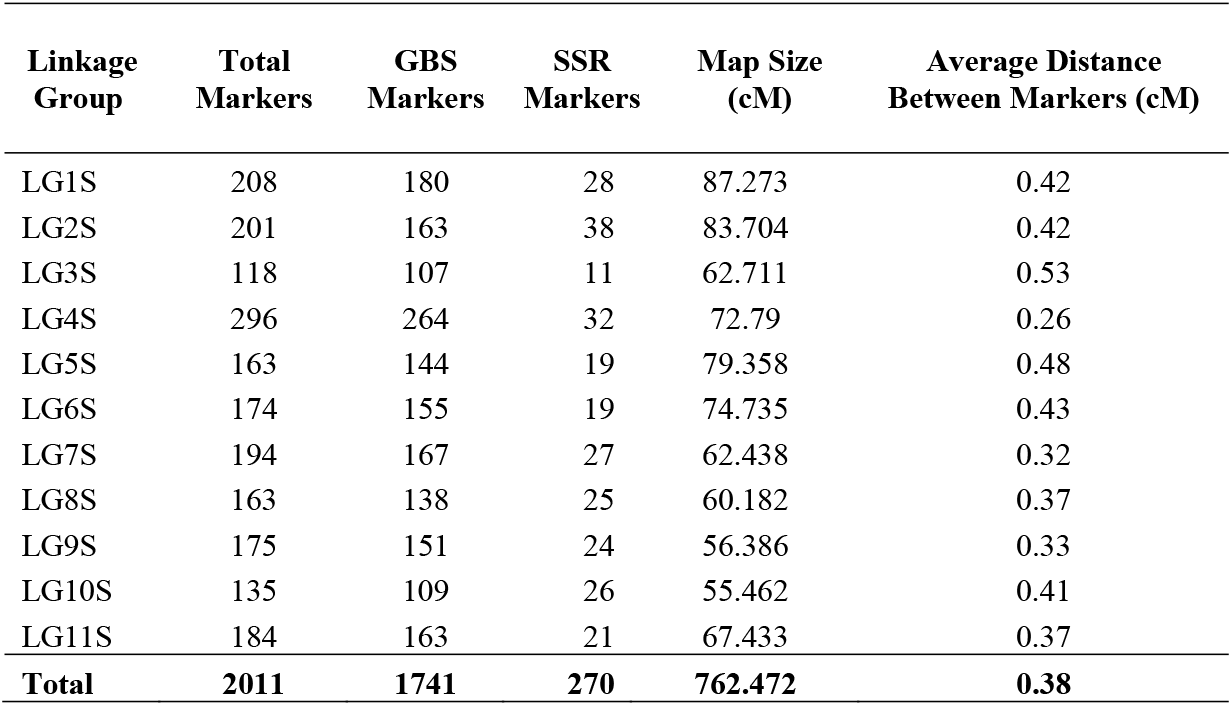
Genetic map statistics of the EFB-susceptible maternal parent (OSU 252.146)

| Linkage Group | Total Markers | GBS Markers | SSR Markers | Map Size (cM) | Average Distance Between Markers (cM) |
| --- | --- | --- | --- | --- | --- |
| LG1S | 208 | 180 | 28 | 87.273 | 0.42 |
| LG2S | 201 | 163 | 38 | 83.704 | 0.42 |
| LG3S | 118 | 107 | 11 | 62.711 | 0.53 |
| LG4S | 296 | 264 | 32 | 72.79 | 0.26 |
| LG5S | 163 | 144 | 19 | 79.358 | 0.48 |
| LG6S | 174 | 155 | 19 | 74.735 | 0.43 |
| LG7S | 194 | 167 | 27 | 62.438 | 0.32 |
| LG8S | 163 | 138 | 25 | 60.182 | 0.37 |
| LG9S | 175 | 151 | 24 | 56.386 | 0.33 |
| LG10S | 135 | 109 | 26 | 55.462 | 0.41 |
| LG11S | 184 | 163 | 21 | 67.433 | 0.37 |
| **Total** | **2011** | **1741** | **270** | **762.472** | **0.38** |

These 11 linkage groups correspond to the haploid chromosome number in hazelnut. Markers are distributed relatively evenly across the map and the number of markers ranges from 118 in LG3 to 296 in LG4. Marker density ranges from an average distance of 0.26 cM in LG4 to 0.53 cM in LG3, and the average distance between markers across the map is 0.38 cM. The paternal genetic map (OSU 414.146) has 1,468 GBS markers and 199 SSR/RAPD markers spanning 908 cM across 10 linkage groups (**Table 2B**). This is one less than the haploid chromosome number suggesting that chromosome corresponding to LG7R merged with other linkage groups. LG7R has low SSR marker density in in previously reported maps [1]. Markers are distributed relatively evenly across the paternal map, and the num-ber of markers ranges from 95 in LG9R to 254 in LG4R. The average distance between adjacent markers is 0.55 cM, and the distances range from 0.63 cM in LG2R to 0.47 cM in LG10R. The EFB-resistant marker, rest-6R, maps to linkage group 6R between SSR marker A12-850TAG-6R and a GBS marker on scaffold C. avellana_Jefferson_10749.

**Table 2B.** Genetic map statistics of the EFB-resistant paternal parent (OSU 414.062)

| Linkage Group | Total Markers | GBS Markers | SSR Markers | Map Size (cM) | Average Distance Between Markers (cM) |
| --- | --- | --- | --- | --- | --- |
| LG1R | 254 | 227 | 27 | 135.146 | 0.53 |
| LG2R | 201 | 175 | 26 | 126.728 | 0.63 |
| LG3R | 178 | 160 | 18 | 85.492 | 0.48 |
| LG4R | 167 | 149 | 18 | 94.656 | 0.56 |
| LG5R | 167 | 146 | 21 | 45.36 | 0.27 |
| LG6R | 150 | 130 | 20 | 84.612 | 0.56 |
| LG8R | 138 | 121 | 17 | 81.735 | 0.59 |
| LG9R | 111 | 95 | 16 | 56.573 | 0.51 |
| LG10R | 180 | 165 | 15 | 84.785 | 0.47 |
| **Total** | **1546** | **1368** | **178** | **795.087** | **0.51** |

The previously reported SSR markers are largely collinear in both maps, verifying mapping accuracy. A total of 1,298 scaffolds from the draft genome assembly are represented in the genetic map, but the proportion of mapped scaffolds is too small to produce a chromosome-scale assembly. The high-density genetic map will be useful for marker-assisted breeding. The full map is available for download on the FTP server.

### Comparative Genomics within the Rosids

Hazelnut belongs to the order Fagales, a large and diverse group with several economically important species including walnut (*Juglandaceae* family), beech and oak (*Fagaceae* family), and birch (*Betulaceae* family). Hazelnut has the first sequenced genome among the Fagales; this map will be useful for comparative genomics in the rosids. Markers from the genetic map were used to assess macro-synteny be-tween hazelnut and peach, which currently has the most complete genome among the rosids (8). A total of 367 markers from the hazelnut genetic map uniquely to peach and the remaining markers either have ambiguous matches or no matches to peach (Fig 2).

**Fig 2.**
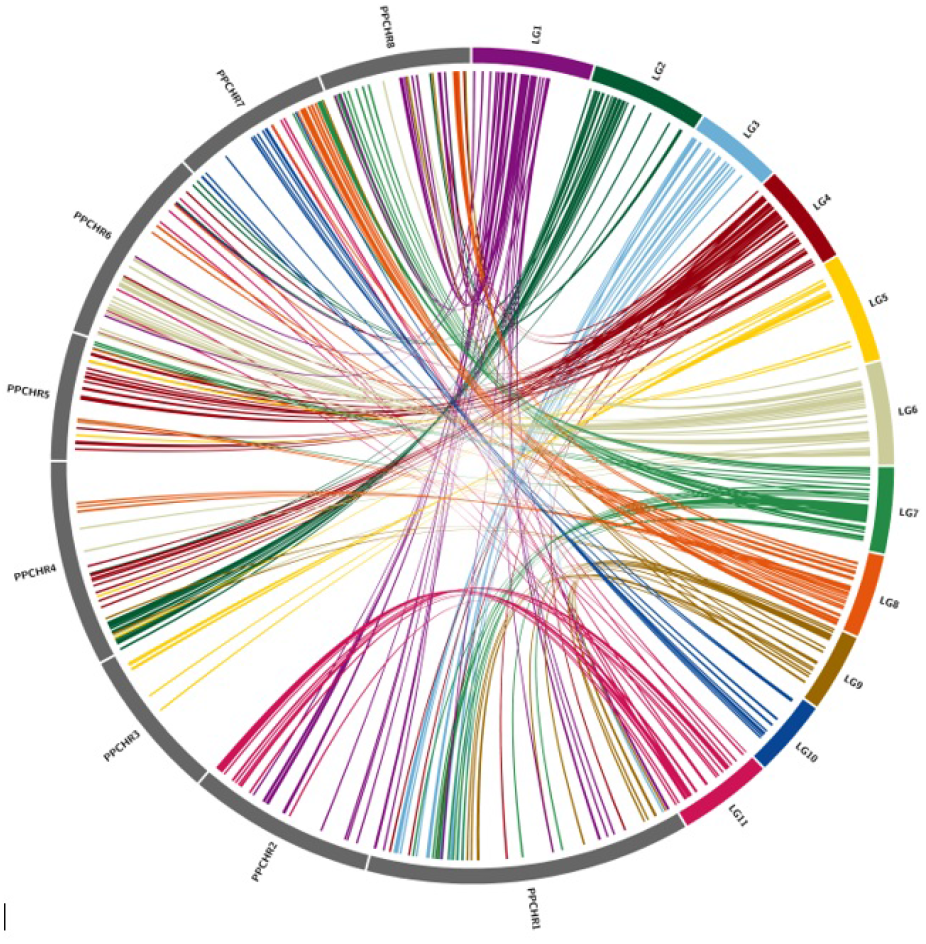
Synteny between the peach and hazelnut genomes based on markers from the genetic map. Links are based on the sequences of flanking markers in the hazelnut genetic map and their homologs in peach. Connections are based on position in the genetic map for hazelnut and physical position in the peach pseudochromosomes

Hazelnut and peach have 1:1 synteny with no evidence of lineage-specific whole-genome duplication in hazelnut. Numerous structural rearrangements are apparent when the peach and hazelnut genomes are compared. Peach chromosome 3 has 1:1 colinearity with LG5, but the remaining chromosomes have a more complex syntenic relationship reflective of chromosome fusions, breakage, and rearrangements that have occurred since the shared ancestral karyotype of nine chromosomes(26). For instance, peach chromosome 1 is collinear to LG3 but also contains regions of LG9 and LG1 from hazelnut.

### Discovery and Characterization of Putative Protein Coding Genes

We have detected protein-coding loci and assigned biological functions to genomic regions of the European hazelnut cultivar ‘Jefferson’. After masking all regions of the genome assembly that were repetitive or low complexity using the program RepeatMasker (10), we implemented the ab initio gene prediction program AUGUSTUS (11) to detect putative protein coding loci within the 36,641 scaffolds and contigs of the draft genome assembly. We trained AUGUSTUS with the known gene features of *Arabidopsis thaliana*; both genera are in the Rosid clade, and the *Arabidopsis* genome is extremely well characterized (12). In total, 36,090 putative coding loci were identified and named Corav_g1.t1 through Corav_g36090.t1.

Sequence homology has been used to assign gene products to putative amino acid sequences in novel de novo assembled genome assemblies; therefore, we aligned the amino acid sequences of the AUGUSTUS output to the NCBI non-redundant protein database using the BLASTP tool (13) with an E-value cut-off of 1*e*^−10^. This homology-based query allowed functional annotation of 23,652 (65.5 percent of the 36,090 putative loci). Of the 23,652 protein sequences mapped to the non-redundant database, 82.5 percent are represented in the well-annotated, phylogenetically related plant genomes *Vitus, Prunus, Populus*, and *Ricinus* (Fig 3).

**Fig 3.**
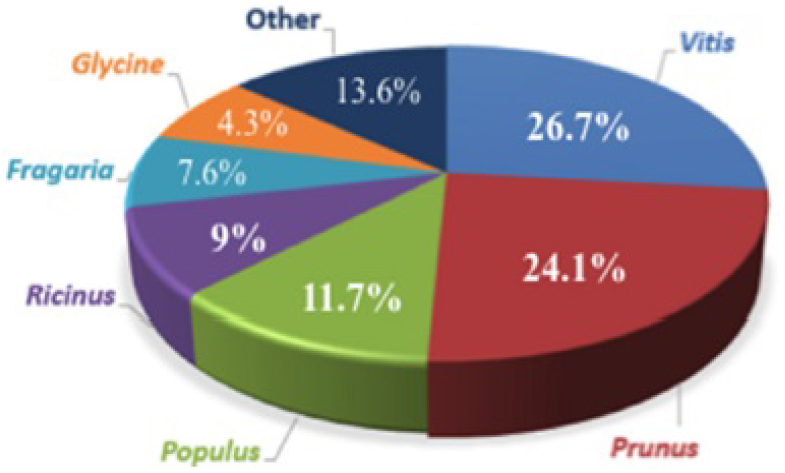
Similarity in protein homology. The majority (82 percent) of protein-coding sequences in the hazelnut assembly are homologous to the related genera *Vitus, Prunus, Populus*, and *Ricinus*.

We then identified all loci annotated as transposable elements, retroelements, and gag-polymerases. This removed 2,398 (1,178 unique) loci from both the annotations and from the putative protein file. These sequences are available for visualization on the hazelnut JBrowse portal as a separate track, and for query on the BLAST portal, both accessed via cavellanagenomeportal.com.

We performed functional classification using gene ontology (GO) analysis to survey, categorize, and define the potential properties of gene products with respect to their predicted biological contexts, using the program BLAST2GO. We were able to functionally classify 11,221 unique loci (31.8 percent of total) comprising 65,536 GO terms, broadly grouped by GO component classifications. Among sequences classified by BLAST2GO, 26.9 percent were assigned to Molecular Function ontology, 18.8 percent were assigned to Cellular Component ontology, and 51 percent were assigned to Biological Process ontology, with only 3 percent lacking assignment to GO classifications.

In total, 22,474 protein-coding regions (64.3 percent of the putative loci) and 2,398 (1,178 unique) transposable elements were identified via homology to the NCBI non-redundant protein database using BLASTP with an E-value cut-off of 1*e*^−10^.

These resources will be useful for the identification of candidate genes underlying traits of interest, for homology-based comparisons to other tree crops, and for the design of molecular markers to enhance breeding efforts. Here we highlight usefulness of the annotations by profiling several interesting gene families with emphasis on their importance to hazelnut breeding.

### Disease-resistance Genes

Plants utilize various defense mechanisms to resist attack by pathogens. Resistance genes, known as R genes, control these mechanisms. R genes confer resistance to specific pathogens, expressing matching avirulence genes in a “gene-for-gene” manner (27). For detailed discussions of the mechanisms of pathogen-host recognition and interactions, readers are directed to recent reviews (28,29). The largest class of resistance genes in plants is the NBS-LRR family (30); the only known functions of the proteins encoded by these genes is in pathogen recognition and defense. The number of NBS-LRR genes varies by species from 57 in cucumber (31) to 200 in *Arabidopsis thaliana* (32) to over 500 in rice (33) and in *Medicago truncatula* (34). There are two functionally distinct subfamilies of NBS-LRR proteins that possessing either a TIR domain or a CC domain upstream of their nucleotide-binding NBS domain. The TIR and CC domains induce distinct downstream response pathways (30).

In the current annotation of the hazelnut genome there are 115 putative NBS-LRR proteins (S1 Table); 35 (30.4 percent) contain TIR-domains, 50 (43 percent) contain CC domains, and 31 (26 percent) have no subfamily designation. In *A. thaliana* the NBS-LRR sequences (and those of most R genes) occur in clusters of closely related sequences around a parent locus (35,36). This is also the case in the hazelnut genome: NBS-LRR genes often occur in clusters as tandem duplicates in the same contig (Fig 4A).

**Fig 4.**
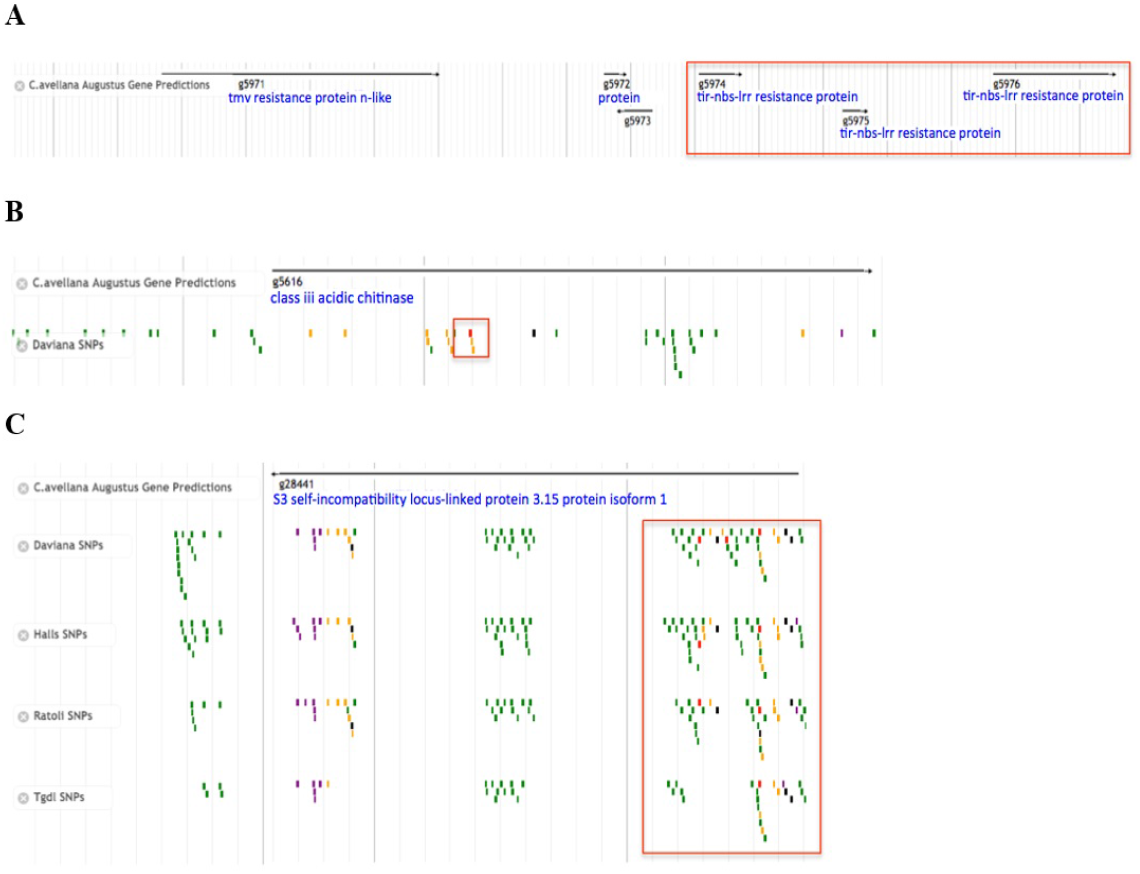
Jbrowse screenshots with added annotation and enhancement displaying: (**A**) Cluster of loci encoding TIR-domain containing NBS-LRR disease resistance proteins, characteristic of many R genes. (**B**) SNP in “putative class III acidic chitinase” in the EFB-susceptible cultivar ‘Daviana’ that introduces a mutation in the splice site acceptor region, possibly increasing susceptibility to fungal attack. (**C**) ‘Daviana’, ‘Hall’s Giant’, ‘Ratoli’, and ‘Tonda Gentile de Langhe’ contain PSC-introducing SNPs in a putative pollen-expressed male determinant SI-related locus.

The annotation of the hazelnut genome assembly will be useful in identifying candidate disease resistance genes for future genetic improvement studies. An example is the ‘Gasaway’ gene, which gives ‘Jefferson’ resistance to EFB. The current annotation is available for BLAST query at hazelnut.blast.mocklerlab.org

### Pollen Incompatibility S-Loci

Hazelnuts are monoecious; separate male and female flowers exist on the same tree. In order to prevent inbreeding and encourage genetic diversity, flowering plants have evolved mechanisms known as self-incompatibility (SI), to promote outcrossing. In dicots, self-incompatibility is inherited as a segregating unit, with pairs of two or more linked genes that encode the male and female incompatibility determinants known as S-haplotypes (37).

The family in which hazelnuts reside, Betulacea, exhibits sporophytic incompatibility (SSI), whereby pollen incompatibility is determined by the parents diploid genotype, inherited as a single locus with multiple alleles (37). There are three genes comprising the core region of the S-locus; the determinant male pollen component has been shown to be a cysteine-rich pollen coat protein (38) encoded by the S-locus cysteine-rich (SCR/SP11). This allele is dominantly expressed in the pollen coat (39). The female determinant is codominant and encoded by two inherited polymorphic loci: the stigma localized glycoprotein (SLG), and the S-locus kinase (SRK).

In angiosperms SI not only provides beneficial self/non-self-determination, it also limits the individuals that may be used in hybrid crosses. For example, if the stigma and pollen express the same allele, SI would render the cross incompatible. This hampers hazelnut-breeding efforts by limiting the parents that can successfully be used for crosses. Fluorescence microscopy is the technique currently used to determine the compatibility of pollinations, and to identify S-alleles in hazelnut cultivars used for crosses (40). To date 33 S-alleles have been identified among hazelnut varieties from several geographical origins (41).

The current ‘Jefferson’ draft genome assembly contains 17 candidates that are annotated as encoding self-incompatibility or S-locus-linked proteins, based on sequence homology to the NCBI nr protein database. Of these 17 loci, 5 are annotated as encoding the stigma localized SLG family (**Table 3A**), while 2 encode the transmembrane-binding SRK (**Table 3B**). The annotation contains 3 loci (**Table 3C**) predicted to encode pollen specific S-locus related proteins, which function as the male determinants. The hazelnut annotation also contains an additional set of 9 loci, which are annotated as S-locus related proteins (**Table 3D**), which may participate in hazelnut SI as well but are currently on different contigs due to the fragmented nature of the assembly.

**Table 3A.** Putative stigma localized glycoprotein (SGL) encoding loci

| Locus | Annotation |
| --- | --- |
| Corav_g6132.t1 | S-locus-specific glycoprotein s6 |
| Corav_g10567.t1 | S-locus-specific glycoprotein s6 |
| Corav_g25571.t1 | S-locus-specific glycoprotein s13 |
| Corav_g34002.t1 | S-locus-specific glycoprotein s6 |
| Corav_g35241.t1 | S-locus-specific glycoprotein s6 |

**Table 3B.** Putative S-locus kinase (SRK) encoding loci

| Locus | Annotation |
| --- | --- |
| Corav_g26148.t1 | S-locus receptor partial |
| Corav_g33469.t1 | S-locus lectin protein kinase family |

**Table 3C.** Putative pollen expressed self-incompatibility loci

| **Locus** | **Annotation** |
| --- | --- |
| Corav_g4284.t1 | S1 self-incompatibility locus-linked pollen protein |
| Corav_g6376.t1 | S3 self-incompatibility locus-linked pollen expressed |
| Corav_g28441.t1 | S3 self-incompatibility locus-linked pollen 3.15 protein isoform 1 |
| Corav_g34002.t1 | S-locus-specific glycoprotein s6 |
| Corav_g35241.t1 | S-locus-specific glycoprotein s6 |

**Table 3D.**
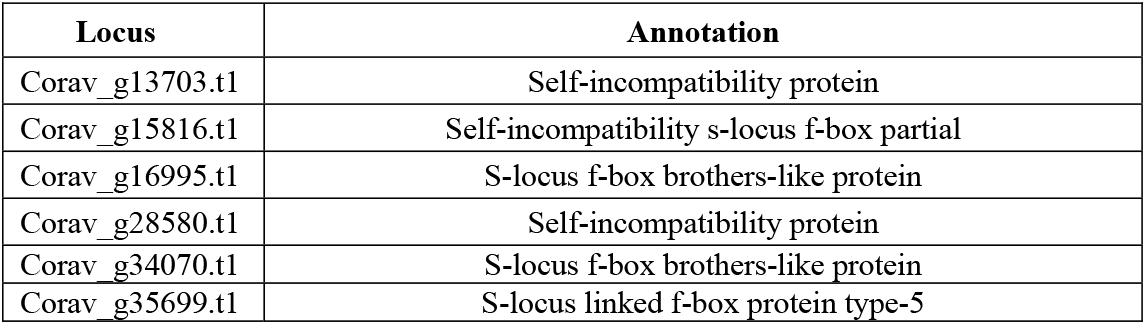
Putative self-incompatibility and related loci

| **Locus** | **Annotation** |
| --- | --- |
| Corav_g13703.t1 | Self-incompatibility protein |
| Corav_g15816.t1 | Self-incompatibility s-locus f-box partial |
| Corav_g16995.t1 | S-locus f-box brothers-like protein |
| Corav_g28580.t1 | Self-incompatibility protein |
| Corav_g34070.t1 | S-locus f-box brothers-like protein |
| Corav_g35699.t1 | S-locus linked f-box protein type-5 |

The identification of the S-alleles that control pollen-stigma incompatibility will assist in the development of PCR primers to quickly screen new hazelnut accessions for compatibility prior to crossing, thus avoiding incompatible crosses, enhance marker assisted selection, identifying the S-locus in hazelnut.

### Resequencing and Polymorphism Detection in Additional Hazelnut Cultivars

An overarching goal in plant breeding is to correlate variations in genomic sequence with agronomic traits of interest. Variations in DNA sequence can occur via singlenucleotide substitutions (SNPs) or by small-scale insertions and deletions (INDELs), ranging in size from a single to hundreds of nucleotides. The vast majority of polymorphisms occur in the less-conserved and non-coding (intergenic) regions rather than protein-coding exonic regions (42). Although these polymorphisms do not affect coding potential, both occurrences have been shown to be under selective pressure and related to agronomic traits (43).

SNPs and INDELs are powerful molecular markers, due in part to their abundance and relative ease of detection in a genome-wide high-throughput experiment (44). Quantitation of SNP and INDEL abundance allows construction of genetic maps and functional markers (44–47). We identified polymorphisms and explored sequence diversity among seven hazelnut cultivars representing four major geo-graphical regions (2,48) of hazelnut production (**Table 4**).

**Table 4.** Resequenced European hazelnut accessions

| **Accession** | **Origin** | **EFB Resistance** | **Description** |
| --- | --- | --- | --- |
| Barcelona | Spain | Intermediate | Important in Oregon |
| Ratoli | Spain | Resistant (Single Gene) | EFB Resistant |
| Daviana | England | Susceptible | Pollinizer for Barcelona |
| Tonda di Giffoni | Italy | Resistant (quantitative) | Excellent kernel quality |
| Tonda Gentile delle Langhe | Italy | Susceptible | Excellent kernel quality |
| Tombul (Extra Ghiaghli) | Black Sea | Susceptible | Important in Turkey |
| Halls Giant | Germany | Intermediate | Pollinizer for Barcelona |

From the Spanish-Italian group: ‘Barcelona’ from Spain; accounting for 60 percent of the hazelnut trees in Oregon and moderately susceptible to EFB, ‘Tonda Gentile delle Langhe’ from northern Italy, with excellent kernel quality and high susceptibility to EFB, ‘Tonda di Giffoni’ from southern Italy, which also has excellent kernel quality and high quantitative resistance to EFB, and ‘Ratoli’ from eastern Spain, which is highly resistant to EFB. ‘Daviana’ represents the English group, a pollinizer for ‘Barcelona’ and highly susceptible to EFB. Representing the Central European group is ‘Halls Giant’, a universal pollinizer which is cold-hardy and moderately resistant to EFB. Finally, from the Black Sea region: ‘Tombul (Extra Ghiaghli)’, a clone of the EFB susceptible Turkish cultivar ‘Tombul’.

### Functional Consequences of Polymorphisms in Protein Coding Loci

The resequencing of seven hazelnut accessions and alignment to the ‘Jefferson’ reference has allowed for the genome-wide association of millions of polymorphisms with the protein coding potential of hundreds of loci. (**Table 5A and 5B**). Additionally, hundreds of protein-coding loci with functions potentially altered or disrupted by the introduction of either a SNP or INDEL were identified (**Table 6A and 6B**).

**Table 5A.**
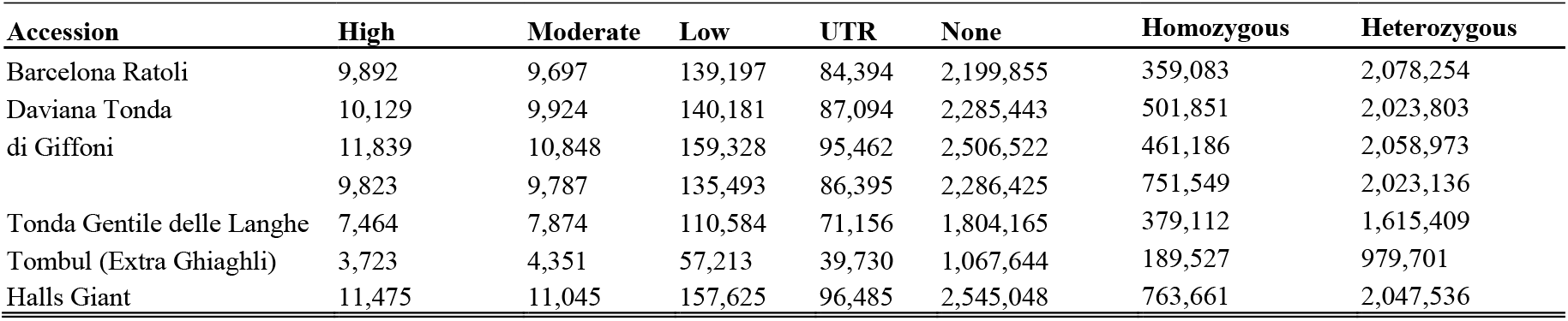
Total SNPs and predicted effect on coding potential between 7 re-sequenced accessions relative to ‘Jefferson’

| **Accession** | **High** | **Moderate** | **Low** | **UTR** | **None** | **Homozygous** | **Heterozygous** |
| --- | --- | --- | --- | --- | --- | --- | --- |
| Barcelona Ratoli | 9,892 | 9,697 | 139,197 | 84,394 | 2,199,855 | 359,083 | 2,078,254 |
| Daviana Tonda | 10,129 | 9,924 | 140,181 | 87,094 | 2,285,443 | 501,851 | 2,023,803 |
| di Giffoni | 11,839 | 10,848 | 159,328 | 95,462 | 2,506,522 | 461,186 | 2,058,973 |
|  | 9,823 | 9,787 | 135,493 | 86,395 | 2,286,425 | 751,549 | 2,023,136 |
| Tonda Gentile delle Langhe | 7,464 | 7,874 | 110,584 | 71,156 | 1,804,165 | 379,112 | 1,615,409 |
| Tombul (Extra Ghiaghli) | 3,723 | 4,351 | 57,213 | 39,730 | 1,067,644 | 189,527 | 979,701 |
| Halls Giant | 11,475 | 11,045 | 157,625 | 96,485 | 2,545,048 | 763,661 | 2,047,536 |

**Table 5B.** Total INDELs and predicted effect on coding potential between 7 re-sequenced accessions relative to ‘Jefferson’

| **Accession** | **High** | **Moderate** | **Low** | **UTR** | **None** | **Homozygous** | **Heterozygous** |
| --- | --- | --- | --- | --- | --- | --- | --- |
| Barcelona Ratoli | 294 | 1,310 | 0 | 15,356 | 321,115 | 45,612 | 285,901 |
| Daviana Tonda | 330 | 1,287 | 0 | 15,801 | 337,077 | 66,222 | 280,171 |
| di Giffoni | 385 | 1,527 | 0 | 17,675 | 375,356 | 102,357 | 281,699 |
|  | 305 | 1,314 | 0 | 15,634 | 336,697 | 58,957 | 285,876 |
| Tonda Gentile delle Langhe | 227 | 1,067 | 0 | 12,530 | 260,172 | 46,596 | 221,003 |
| Tombul (Extra Ghiaghli) | 117 | 596 | 0 | 6,845 | 154,162 | 24,768 | 134,123 |
| Halls Giant | 354 | 1,530 | 0 | 17,812 | 379,194 | 98,933 | 287,029 |

**Table 6A.**
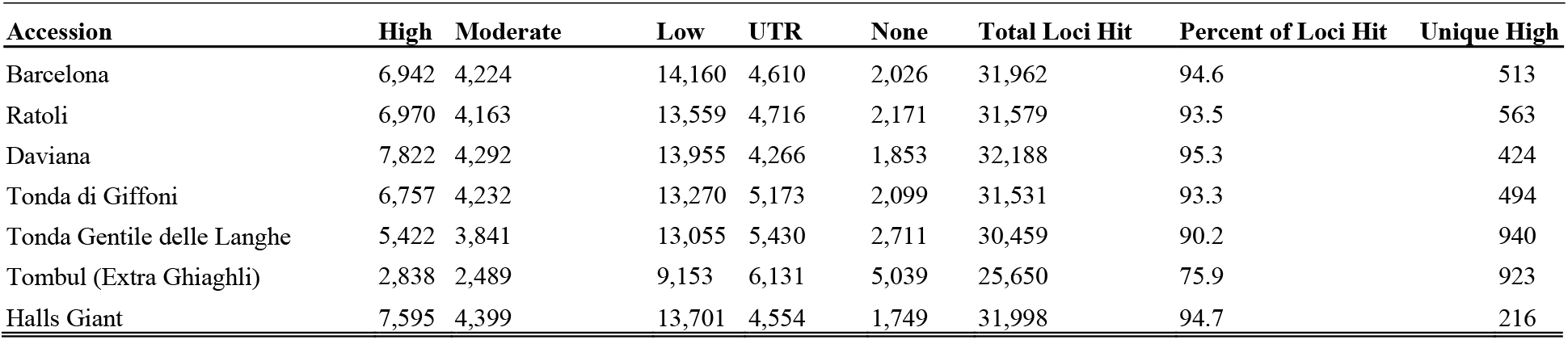
Unique loci containing SNPs and predicted effect on coding potential between re-sequenced accessions relative to ‘Jefferson’

| **Accession** | **High** | **Moderate** | **Low** | **UTR** | **None** | **Total Loci Hit** | **Percent of Loci Hit** | **Unique High** |
| --- | --- | --- | --- | --- | --- | --- | --- | --- |
| Barcelona | 6,942 | 4,224 | 14,160 | 4,610 | 2,026 | 31,962 | 94.6 | 513 |
| Ratoli | 6,970 | 4,163 | 13,559 | 4,716 | 2,171 | 31,579 | 93.5 | 563 |
| Daviana | 7,822 | 4,292 | 13,955 | 4,266 | 1,853 | 32,188 | 95.3 | 424 |
| Tonda di Giffoni | 6,757 | 4,232 | 13,270 | 5,173 | 2,099 | 31,531 | 93.3 | 494 |
| Tonda Gentile delle Langhe | 5,422 | 3,841 | 13,055 | 5,430 | 2,711 | 30,459 | 90.2 | 940 |
| Tombul (Extra Ghiaghli) | 2,838 | 2,489 | 9,153 | 6,131 | 5,039 | 25,650 | 75.9 | 923 |
| Halls Giant | 7,595 | 4,399 | 13,701 | 4,554 | 1,749 | 31,998 | 94.7 | 216 |

**Table 6B.** Unique loci containing INDELs and predicted effect on coding potential between re-sequenced accessions relative to ‘Jefferson’

| **Accession** | **High** | **Moderate** | **Low** | **UTR** | **None** | **Total Loci Hit** | **Percent of Loci Hit** | **Unique High** |
| --- | --- | --- | --- | --- | --- | --- | --- | --- |
| Barcelona | 285 | 1,198 | 0 | 9,845 | 10,857 | 22,185 | 65.7 | 67 |
| Ratoli | 321 | 1,177 | 0 | 10,015 | 10,728 | 22,241 | 65.8 | 83 |
| Daviana | 371 | 1,374 | 0 | 10,823 | 10,706 | 23,274 | 68.9 | 58 |
| Tonda di Giffoni | 299 | 1,205 | 0 | 9,958 | 10,653 | 22,115 | 65.5 | 79 |
| Tonda Gentile delle Langhe | 223 | 988 | 0 | 8,364 | 10,610 | 20,185 | 59.8 | 126 |
| Tombul (Extra Ghiaghli) | 115 | 569 | 0 | 4,963 | 9,071 | 14,718 | 43.6 | 124 |
| Halls Giant | 346 | 1,388 | 0 | 10,824 | 10,635 | 23,193 | 68.7 | 34 |

The discussion of several of these polymorphisms seeks to underscore the value of these predictions in candidate discovery and hypothesis generation for future experiments. The complete annotated variant files for each accession used in this study are hosted on the FTP server at hazelnut.data.mocklerlab.org, available for both download and query.

Each of the seven resequenced accessions contains dozens of polymorphisms in putative disease resistance genes; here we focus on annotated variants unique to each accession. The English accession ‘Daviana’, which is susceptible to EFB, has a SNP in the “putative class III acidic chitinase” encoding locus Corav_g5616.t1 (**Fig 4B**). Plant chitinases play a role in defense from fungal pathogens by interrupting vegetative growth of fungal hyphae (49). Overexpression of plant chitinases, often in combination with PR proteins (50), have led to improved fungal resistance in many crop systems. The polymorphism in locus Corav_g5616.t1 introduces a mutation into a splice site acceptor region relative to contig C.avellana_Jefferson_00575; this may result in a mis-spliced transcript destined for degradation prior to translation. It is possible that these variants in resistance genes increase the susceptibility of ‘Daviana’ to fungal attack, such as infection with EFB.

Three of the resequenced accessions are fully susceptible to EFB: the English accession ‘Daviana’, the Italian accession ‘Tonda Gentile delle Langhe’, and the Turkish accession ‘Tombul (Extra Ghiaghli)’. Uniquely shared among these accessions are 12 loci containing SNPs that introduce functionally deleterious HIGH effects. Two of these loci are known disease resistance genes: Corav_g5237.t1, a TIR-NBS-LRR resistance protein and Corav_g34429.t1, a member of the “blight-associated protein p12 precursor” proteins. Two of the five members of this family in the hazelnut annotation have polymorphisms in the EFB-susceptible accessions that are predicted to alter coding potential.

Perhaps one of the most interesting groups of genes with annotated variants is that of the putative SI genes. Of the 17 loci in the hazelnut annotation annotated as participating in SI, five of the SSI related loci have SNPs that are highly likely to impact protein production (**Table 7**).

**Table 8.** Variants in S-loci annotated as affecting coding potential between re-sequenced accessions

| **Locus** | **Annotation** | **Barcelona** | **Daviana** | **Halls Giant** | **Ratoli** | **Tonda GdL** | **Tombul** | **Tonda di Giffoni** |
| --- | --- | --- | --- | --- | --- | --- | --- | --- |
| Corav_g6132.t1 | S-locus-specific glycoprotein s6 | SNP | SNP | SNP | SNP | SNP | SNP | SNP |
| Corav_g6376.t1 | S3 self-incompatibility locus-linked pollen expressed | SNP | SNP | SNP | SNP | SNP | SNP | SNP |
| Corav_g10567.t1 | S-locus-specific glycoprotein s6 | SNP | SNP | SNP |  |  | SNP |  |
| Corav_g28441.t1 | S3 self-incompatibility locus-linked pollen 3.15 protein isoform 1 |  | SNP | SNP | SNP | SNP | SNP |  |

These variants may underlie the genetic SI mechanism, which leads to the incompatibility phenotype observed in hazelnut. For example, Corav_g28441.t1, a putative male determinant “S3 self-incompatibility locus-linked pollen 3.15 protein isoform 1” encoding locus, has SNPs in the accessions ‘Daviana’, ‘Halls Giant’, ‘Ratoli’, and Tonda Gentile de Langhe’ (**Fig 4C**), leading to a premature stop codon (PSC).

## Conclusions

The association of genes with desirable traits has long been a goal of plant breeding, and having an annotated and characterized genome assembly is the first step in realizing this goal. We have sequenced, assembled, and characterized the draft genome of the European hazelnut (*Corylus avellana* L.) accession ‘Jefferson’ and resequenced seven additional cultivars, Hazelnut is the first sequenced plant in the order Fagales, which includes walnut, pecan and several economically important wood species including oak, beech and hickory. Gene prediction and functional annotation of protein-coding loci allowed identification of agriculturally relevant loci and will be extremely useful for future molecular characterization and marker discovery. Variations in DNA sequence between ‘Jefferson’ and resequenced accessions range in size from single nucleotide polymorphisms to hundreds of nucleotides. Predicting the effects of these variations on the coding potential of gene loci are an integral step in the identification of genes underlying traits of interest and hypothesis generation for future molecular experiments. The establishment of PCR markers corresponding to genes of interest, such as those controlling the pollen-stigma incompatibility, can be used to rapidly screen additional accessions to determine the presence of the gene of interest prior to using as parents in breeding experiments. In addition, one these markers are established, the progeny of these crosses can be screened via PCR to determine whether they possess the desired trait.

Disease resistance is often involves several loci and pathways; the resistance and susceptibility to EFB displayed in certain accessions (**Table 4**) is highly unlikely to involve the same genes. For example, the EFB resistance locus in the Spanish accession ‘Ratoli’ maps to a different linkage group than that of the cultivar ‘Gasaway’ and may offer a more robust source of resistance (22,51). Variation in nucleotide sequences among hazelnut populations in different geographic regions, have the potential to introduce polymorphisms in coding loci. These polymorphisms may affect the susceptibility of these cultivars to disease. The establishment of ‘Jefferson’ as the reference hazelnut cultivar will aid in unraveling the genes responsible for disease resistance among these cultivars. Once new sources of resistance are identified, it will be possible to further enhance resistance by stacking the traits of multiple accessions through several rounds of selective breeding. It may be possible to utilize wild accessions as parents in crosses; the discovery of new sources of EFB resistance in American hazelnut would allow the creating of hybrid cultivars with European hazel.

As ‘Jefferson’ is the F1 progeny of a clonal cross, it contains heterozygous regions within its genome, as demonstrated by the bimodal K-mer distribution. This contributed to the fragmented nature of the assembly; the short-read sequencing technology and programs used for genome assembly were unable to resolve the more heterozygous regions of the genome, resulting in breaks in the assembly and generation of many smaller contigs. A portion of the assembly may also exist as mosaics, full resolution of which will be realized as the draft assembly is improved with future sequencing experiments.

Despite the fragmented contigs, the initial de novo assembly of the hazelnut genome captures over 90 percent of the empirically determined hazelnut genome size, and has been used to identify hundreds of new SSR markers (S.A. Mehlenbacher, personal communication, April 2016). The current draft assembly will be useful for the alignment of future physical markers and sequenced BAC libraries, the identification of additional molecular markers, and alignment of RNA-seq reads from experiments that, for example, survey differential gene expression from incompatible and compatible pollinations over time.

At the writing of this manuscript, the ‘Gasaway’ allele conferring resistance to EFB has not yet been identified. However, Sathuvalli *et al*. (52,53) generated BAC libraries and identified polymorphic RAPD markers linked to EFB resistance. Alignment of these BAC end markers to the ‘Jefferson’ assembly resulted in matches of 100 percent identify across their length (data not shown), but the fragmented nature of the assembly and small contigs did not allow for the full resolution of the downstream sequence nor positive identification of the gene. This is not surprising given the fact markers may be located megabases away from the loci they segregate with; such resolution is not possible given the current fragmented assembly.

GBS analysis of the hazelnut F1 mapping population resulting from the cross of the EFB-susceptible maternal parent OSU 252.146 and the resistant paternal parent 414.062 added an additional 3,209 markers to the existing genetic linkage map (22). This improved quality genetic linkage map represents a 5-fold improvement over the existing map. The average distance between markers in the current map is 0.38 cM. This resolution allows genome-wide analyses of molecular variation and additional marker discovery to accelerate molecular breeding in hazelnut. For example, the EFB-resistance in ‘Gasaway’ was previously assigned to LG6 (22), and the separate gene responsible for resistance in the Spanish cultivar ‘Ratoli’ segregates on LG7 (51). In addition, the recently characterized accession OSU 759.010 transmits resistance via a locus on LG2 (53) different from locations of other identified resistance loci. The variants discovered and characterized in this study, including the new GBS-derived markers placed on the genetic linkage map, will be useful for identifying candidate genes responsible for EFB resistance and other genetic factors that reduce yield. This research will also complement a recently released genetic linkage map, constructed from the progeny of a cross between the cultivars ‘Tonda Gentile delle Langhe’ x ‘Halls Giant’, developed to identify QTLs for traits such as pest tolerance and nut quality in Italian cultivars (54).

The ‘Jefferson’ reference hazelnut genome, annotations, and other resources are available to the public on https://cavellanagenomeportal.com. An interactive Jbrowse interface allows visualization of the annotations, alignments, and polymorphisms f or each accession in separate overlayable tracks. Variant calls for each of the seven resequenced accessions annotated with their putative effects on coding potential and a BLAST portal increase the available genomic resources for European hazelnut.

## Supporting information

## Author’s contributions

ERR and SAM collected the hazelnut tissues. ERR extracted the DNA. DWB conducted the sequence assembly. ERR conducted the sequence analysis and wrote the manuscript. RV conducted the GBS analysis and wrote the manuscript. HDP developed the web interfaces and conducted the data hosting. SAM and TCM conceived of the study, participated in its design and coordination, and provided funding. All authors read and approved the final manuscript.

## Acknowledgements

We would like to thank the Georgia Genomics Facility at the University of Georgia for the preparation of Illumina libraries; Anne-Marie Girard, Caprice Rosato, Mark Dasenko, and Mathew Peterson for qualitative assessment of the libraries, Illumina cluster generation, and computational support (Center for Genome Research and Biocomputing, Oregon State University); scientists at MOgene (St. Louis) for construction and sequencing of the GBS libraries. This work was supported by the Oregon State University Agricultural Research Foundation, the Oregon Hazelnut Commission, and the Donald Danforth Plant Science Center.

## Supplemental Materials

Supplementary Table 1

