## Supplementary material for "A Draft Genome and High-Density Genetic Map of European Hazelnut (*Corylus avellana* L.)"

**Supplementary Table 1.** Loci annotated as encoding members of the NBS-LRR family of disease resistance proteins

| Locus | Annotation |
|---|---|
| Corav_g10139.t1 | nbs-lrr resistance protein |
| Corav_g10150.t1 | cc-nbs-lrr resistance protein |
| Corav_g11411.t1 | cc-nbs-lrr resistance protein |
| Corav_g11649.t1 | tir-nbs-lrr resistance protein |
| Corav_g11670.t1 | cc-nbs-lrr resistance protein |
| Corav_g11774.t1 | nbs-lrr resistance protein |
| Corav_g12203.t1 | cc-nbs-lrr resistance protein |
| Corav_g12647.t1 | nbs-lrr resistance protein |
| Corav_g12810.t1 | nbs-lrr resistance protein |
| Corav_g13195.t1 | tir-nbs-lrr type disease resistance protein |
| Corav_g13212.t1 | tir-nbs-lrr resistance protein |
| Corav_g14247.t1 | cc-nbs-lrr resistance protein |
| Corav_g14428.t1 | nbs-lrr resistance protein |
| Corav_g14499.t1 | tir-nbs-lrr resistance protein |
| Corav_g15327.t1 | cc-nbs-lrr resistance protein |
| Corav_g15370.t1 | cc-nbs-lrr resistance protein |
| Corav_g15789.t1 | nbs-lrr resistance protein |
| Corav_g16063.t1 | tir-nbs-lrr resistance protein |
| Corav_g16645.t1 | tir-nbs-lrr resistance protein |
| Corav_g16648.t1 | tir-nbs-lrr resistance protein |
| Corav_g17256.t1 | cc-nbs-lrr resistance protein |
| Corav_g17343.t1 | nbs-lrr resistance protein |
| Corav_g17354.t1 | cc-nbs-lrr resistance protein |
| Corav_g17662.t1 | cc-nbs-lrr resistance protein |
| Corav_g17664.t1 | cc-nbs-lrr resistance protein |
| Corav_g17912.t1 | nbs-lrr disease resistance protein |
| Corav_g17995.t1 | tir-nbs-lrr resistance protein |
| Corav_g18241.t1 | tir-nbs-lrr resistance protein |
| Corav_g18437.t1 | cc-nbs-lrr resistance protein |
| Corav_g18438.t1 | cc-nbs-lrr resistance protein |
| Corav_g18823.t1 | cc-nbs-lrr resistance protein |

| | |
|---|---|
| Corav_g19051.t1 | nbs-lrr resistance protein |
| Corav_g19177.t1 | cc-nbs-lrr resistance protein |
| Corav_g19555.t1 | tir-nbs-lrr rct1-like resistance protein |
| Corav_g1961.t1 | cc-nbs-lrr resistance protein |
| Corav_g1964.t1 | cc-nbs-lrr resistance protein |
| Corav_g19735.t1 | cc-nbs-lrr resistance protein |
| Corav_g19839.t1 | cc-nbs-lrr resistance protein |
| Corav_g2236.t1 | tir-nbs-lrr resistance protein |
| Corav_g25042.t1 | tir-nbs-lrr resistance protein |
| Corav_g25082.t1 | nbs-lrr resistance protein |
| Corav_g25083.t1 | cc-nbs-lrr resistance protein |
| Corav_g25256.t1 | cc-nbs-lrr resistance protein |
| Corav_g25488.t1 | nbs-lrr disease-resistance protein scn3r1 |
| Corav_g25503.t1 | cc-nbs-lrr resistance protein |
| Corav_g25858.t1 | tir-nbs-lrr resistance protein |
| Corav_g25875.t1 | tir-nbs-lrr resistance protein |
| Corav_g26332.t1 | cc-nbs-lrr resistance protein |
| Corav_g26374.t1 | cc-nbs-lrr resistance protein |
| Corav_g26423.t1 | nbs-lrr resistance protein |
| Corav_g26430.t1 | cc-nbs-lrr resistance protein |
| Corav_g26460.t1 | tir-nbs-lrr type disease resistance protein |
| Corav_g26578.t1 | tir-nbs-lrr resistance protein |
| Corav_g26696.t1 | cc-nbs-lrr resistance protein |
| Corav_g26805.t1 | cc-nbs-lrr resistance protein |
| Corav_g26808.t1 | cc-nbs-lrr resistance protein |
| Corav_g27103.t1 | tir-nbs-lrr type disease resistance protein |
| Corav_g27126.t1 | nbs-lrr disease-resistance protein scn3r1 |
| Corav_g27512.t1 | cc-nbs-lrr resistance protein |
| Corav_g27755.t1 | nbs-lrr disease-resistance protein scn3r1 |
| Corav_g27780.t1 | cc-nbs-lrr resistance protein |
| Corav_g27965.t1 | nbs-lrr resistance protein |
| Corav_g28151.t1 | cc-nbs-lrr resistance protein |
| Corav_g28392.t1 | cc-nbs-lrr resistance protein |
| Corav_g28411.t1 | cc-nbs-lrr resistance protein |
| Corav_g28500.t1 | nbs-lrr resistance protein |

| | |
|---|---|
| Corav_g28508.t1 | nbs-lrr type disease resistance-like partial |
| Corav_g28728.t1 | bed finger-nbs-lrr resistance protein |
| Corav_g28782.t1 | tir-nbs-lrr resistance protein |
| Corav_g29384.t1 | tir-nbs-lrr resistance protein |
| Corav_g29600.t1 | nbs-lrr disease-resistance protein scn3r1 |
| Corav_g29671.t1 | nbs-lrr resistance protein |
| Corav_g30297.t1 | cc-nbs-lrr resistance protein |
| Corav_g30674.t1 | cc-nbs-lrr resistance protein |
| Corav_g30701.t1 | nbs-lrr class disease resistance protein |
| Corav_g30834.t1 | cc-nbs-lrr resistance protein |
| Corav_g31136.t1 | cc-nbs-lrr resistance protein |
| Corav_g31235.t1 | nbs-lrr resistance protein |
| Corav_g31257.t1 | nbs-lrr resistance protein |
| Corav_g31348.t1 | tir-nbs-lrr disease resistance protein |
| Corav_g31479.t1 | tir-nbs-lrr resistance protein |
| Corav_g31575.t1 | nbs-lrr disease-resistance protein scn3r1 |
| Corav_g31703.t1 | cc-nbs-lrr resistance protein |
| Corav_g32104.t1 | tir-nbs-lrr resistance protein |
| Corav_g32463.t1 | tir-nbs-lrr resistance protein |
| Corav_g32562.t1 | cc-nbs-lrr resistance protein |
| Corav_g32830.t1 | cc-nbs-lrr resistance protein |
| Corav_g32873.t1 | cc-nbs-lrr resistance protein |
| Corav_g32888.t1 | cc-nbs-lrr resistance protein |
| Corav_g32944.t1 | cc-nbs-lrr resistance protein |
| Corav_g32951.t1 | tir-nbs-lrr resistance protein |
| Corav_g3297.t1 | tir-nbs-lrr resistance protein |
| Corav_g32971.t1 | cc-nbs-lrr resistance protein |
| Corav_g33056.t1 | cc-nbs-lrr resistance protein |
| Corav_g33192.t1 | nbs-lrr resistance protein |
| Corav_g33224.t1 | cc-nbs-lrr resistance protein |
| Corav_g34032.t1 | nbs-lrr resistance protein |
| Corav_g34214.t1 | tir-nbs-lrr resistance protein |
| Corav_g34390.t1 | tir-nbs-lrr resistance protein |
| Corav_g35374.t1 | cc-nbs-lrr resistance protein |

| | |
|---|---|
| Corav_g35520.t1 | tir-nbs-lrr resistance protein |
| Corav_g35541.t1 | cc-nbs-lrr resistance protein |
| Corav_g35820.t1 | cc-nbs-lrr resistance protein |
| Corav_g36019.t1 | cc-nbs-lrr resistance protein |
| Corav_g5122.t1 | nbs-lrr resistance protein |
| Corav_g5237.t1 | tir-nbs-lrr resistance protein |
| Corav_g5587.t1 | tir-nbs-lrr resistance protein |
| Corav_g5589.t1 | nbs-lrr resistance protein |
| Corav_g5974.t1 | tir-nbs-lrr resistance protein |
| Corav_g5975.t1 | tir-nbs-lrr resistance protein |
| Corav_g5976.t1 | tir-nbs-lrr resistance protein |
| Corav_g8198.t1 | tir-nbs-lrr resistance protein |
| Corav_g840.t1 | nbs-lrr resistance protein |
| Corav_g841.t1 | nbs-lrr resistance protein |
| Corav_g904.t1 | tir-nbs-lrr resistance protein |
